# OFF-periods reduce the complexity of neocortical activity during sleep

**DOI:** 10.1101/2021.06.11.448131

**Authors:** Joaquín González, Matias Cavelli, Adriano BL Tort, Pablo Torterolo, Nicolás Rubido

## Abstract

Complexity of electroencephalographic signals decreases during slow-wave sleep (SWS), however, the neural mechanism for this decrease remains elusive. Here, we show that this complexity reduction is caused by synchronous neuronal OFF-periods. We analyse *in-vivo* recordings from neocortical neuronal populations, finding that OFF-periods in SWS trap cortical dynamics, making the population activity more recurrent, deterministic, and less chaotic than during REM sleep or Wakefulness. Importantly, when we exclude OFF-periods, SWS becomes indistinguishable from Wakefulness or REM sleep. In fact, we show that spiking activity for all states has a universal scaling compatible with critical phenomena. We complement these results by a critical branching model that replicates our experimental findings, where we show that forcing OFF-periods to a percentage of neurons suffices to generate a decrease in complexity that replicates SWS.

## Introduction

Complexity in brain dynamics has been suggested as a necesary condition for consciousness [1–3]. Accordingly, the complexity of electroencephalographic (EEG) signals is observed to decrease during unconscious states, such as during sleep [4–13] or anesthesia [14–20]. Generally, the complexity of a signal refers to its diversity of patterns, which can be estimated from different complexty or information metrics – as seen from the increasing number of publications over the last decade [1–20]. But in spite of the widely observed decrease in complexity of field recordings during unconscious states, we lack a description to the neural mechanism causing it; with some reports even challenging its relevance for consciousness [21].

In contrast, *in vitro* cortical populations show complex patterns of spontaneus spiking [22, 23], which simplify during the occurrence of induced slow-waves [24]. Similar slow-waves characterize slow-wave sleep (SWS), originating from synchronous periods of spiking silences, i.e., OFF-periods [25–29]. These OFF-periods reflect hyperpolarized (Down) states of neocortical neurons [30] that disrupt neural intereactions [31], pointing to a neural mechanism that could explain the decrease in complexity of field recordings. For example, a sleep-like OFF-period has been reported to affect complexity in patients with unresponsive wakefulness syndrome [32]. Nevertheless, the hypothesis that OFF-periods reduce the complexity of cortical activity during sleep is yet to be demonstrated.

Here, we study *in-vivo* neuronal recordings from the neocortex and hippocampus of 15 rats cycling through the states of wakefulness (Wake), slow-wave sleep (SWS), and rapid-eye movement (REM) sleep. These spike records total ≃ 1600 neurons over 31 sessions (each session measures 51 ± 5 neurons) – hereafter referred as population activity. Our analyses show that the population activity is less complex during SWS because it gets periodically trapped into recurrent, deterministic, non chaootic states during OFF-periods. In addition, we show that spiking (active) periods during SWS are indistinguishable from those of Wake or REM sleep, following a universal scaling similar to those found in critical phenomena. We confirm these observations by modelling the cortex as a critical branching process, which reproduces our *in-vivo* results and shows that the introduction of OFF-periods is sufficient to reproduce a SWS-like state. Overall, our results indicate that the decrease in complexity is a sleep trait stemming from the presence of synchronous OFF-periods in a near-critical system.

## Results

### Recurrence analysis reduces high-dimensional dynamics to a 2*D* representation

The population activity (details in Datasets) from a cortical location at any given time is a high-dimensional variable detailing the system’s instantaneous state (Fig. 1**a**). Its evolution gives a trajectory, which has the firing counts of each neuron as they evolve through phase-space (Fig. 1**b**). An attractor is evidenced as a manifold that attracts different trajectories of the system to the same region of phase-space – the more convoluted (fractal) the attractor is, the higher the temporal complexity of its trajectories. However, the attractor resulting from the trajectory of any given cortical area is typically high-dimensional; note that 50 neurons hold a ~ 50*D* phase-space. By applying Recurrence Quantification Analysis (RQA) we reduce these dimensions to the analysis of 2*D* recurrence plots (RP) (Fig. 1**c**).

**Figure 1:**
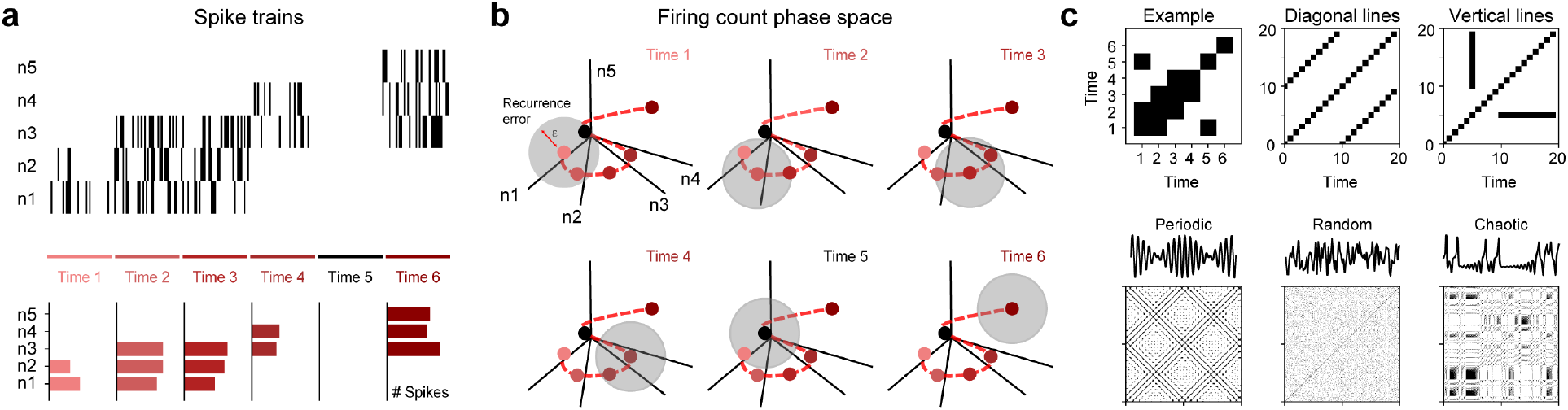
Recurrence example of population activity. **a** Example of spike trains for 5 neurons (n1-n5). Their firing counts are shown in the bottom panel for the respective time-bins, where an OFF-period can be seen at Time 5. **b** Resultant phase-space trajectory (evolution), where axes are the firing count variables of each neuron – trajectory is shown by a dashed line and (binned) measurements by filled circles. For each measurement, a ball of radius *ϵ* (shaded grey area) is used to find when the trajectory recurs to the same region, defining a recurrence plot (RP). **c** (Top left panel) RP for trajectory from **b**. (Remaining panels) RP examples, showing diagonal lines, vertical lines, as well as periodic, random, and chaotic trajectories.

We construct RP as follows. Let 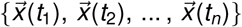 be a trajectory, where 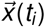 is the state-vector whose components are firing counts, *x_k_*(*t_i_*) (*k* = 1, …, *N*), for each neuron in the population at time, *t_i_*, with *i* = 1, …, *T*, *T* being the length of the recording; set to *T* =10 *s*. We find *x_k_* by adding the *k*-th neuron spike-trains within 50 *ms* time-bins; set to *t*_*i*+1_ – *t_i_* = 50 *ms* ∀ *i*. We choose this time-bin width to match the definition of an OFF-period, i.e., a period ≥ 50 *ms* without spikes. Hence, our firing counts are integer variables that can range from 0 up to 50 (assuming a maximum of 1 spike per *ms*). An RP is then defined by a symmetric matrix whose entries are: *R*(*i, j*) = 1 if 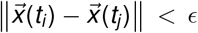, or *R*(*i*, *j*) = 0 otherwise, with *i*, *j* = 1, …, *T* and *ϵ* > 0 defining closeness. A recurrence happens whenever the system’s trajectory returns to the same region of phase-space up to *ϵ*. We set *ϵ* = *σ_p_* (*σ_p_* being the standard deviation of the population-activity during wakefulness) to guarantee a sufficiently sparse RP but with sufficient points to carry statistical analyses. Nevertheless, our results are robust to changes in *ϵ* or time-bin width (Fig. S1).

Two generic structures appear in RP: diagonal lines, originating from periodic trajectories, and vertical lines, originating from trapped (frozen) trajectories. These structures help to differentiate between periodic, random, or chaotic trajectories (corresponding panels in Fig. 1**c**), which can be quantified by different metrics (see RQA in Methods). We use RQA to measure (i) Recurrence Rates (density of points in a RP), RR, (ii) Determinism (proportion of points forming diagonal lines), DET, (iii) Laminarity (proportion of points forming vertical lines), LAM, (iv) Trapping Times (average length of vertical lines), TT, and (v) Divergences (inverse of the longest diagonal line), DIV. RR quantifies the overall recurrence of the system, DET measures the smoothness of trajectories, LAM quantifies the proportion of recurrences caused by trapped states, TT measures the average time the system spends in a trapped state, and DIV quantifies the chaoticity of the trajectory. Thus, predictability in the system’s trajectory is quantified by RR, DET, LAM, and TT, where the larger [smaller] their values, the more [less] predictable. On the other hand, chaoticity is quantified by DIV, where the larger [smaller] its value the more divergent [convergent] is the trajectory.

### Neuronal activity decreases its complexity during slow-wave sleep

Here we show RQA results for the frontal cortex (~ 900 neurons) during Wake, SWS, and REM sleep. Figure 2**a** shows a representative local field potential (LFP) and the spike-trains of the frontal-cortex neurons in a session (Neuron #). The recurrence plots (RP) for each state are shown in Fig. 2**b**, which are constructed from 10 s window trajectories of firing counts. From these panels, we note that SWS exhibits a denser RP than Wake or REM sleep; namely, SWS has firing patterns that recur more often than Wake or REM sleep. Also, SWS shows a distinctive square-shaped recurrence-pattern lasting approximately 100 *ms*, which points to a periodic trajectory with trapped states. RQA confirms that frontal-cortex activity during SWS is significantly more predictable and less chaotic than Wake or REM sleep, i.e., less complex (Fig. 2**c**) – statistics are shown in Table S1. Specifically, SWS has the largest RR, DET, LAM and TT, indicating a higher predictability for the neuronal activity during SWS than during Wake or REM sleep, whereas DIV is larger during Wake and REM sleep than during SWS, indicating that SWS is significantly less chaotic.

**Figure 2:**
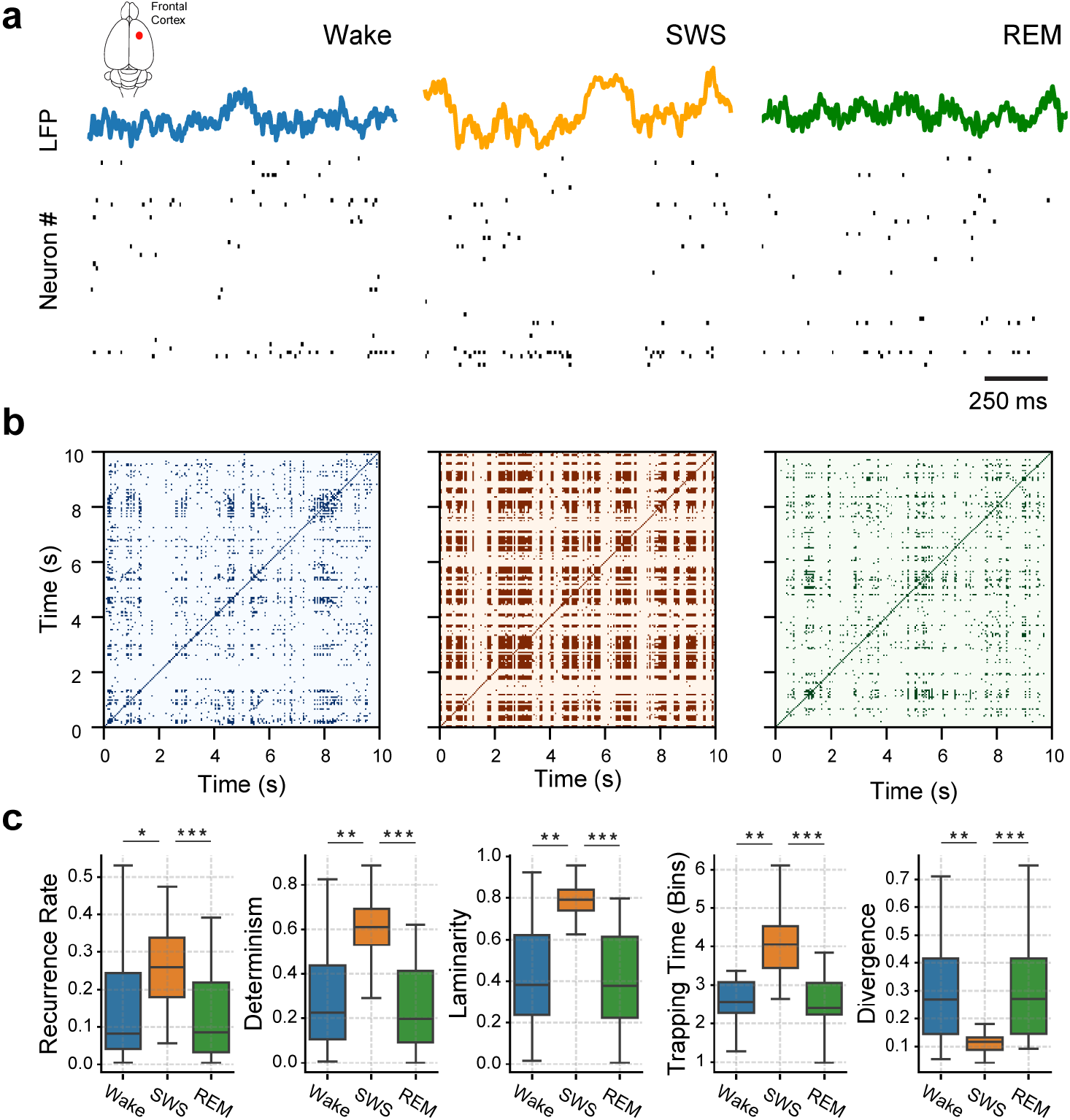
Recurrence analysis of *in-vivo* population activity from the frontal cortex. **a** Local field potentials (top trace) and spike-train raster plots (1 *s* interval) for a representative rat during Wake (left panel), SWS (middle panel), and REM sleep (right panel). **b** Respective recurrence plots constructed from a 10 *s* interval of the population activity after binning the spike trains into 50 *ms* windows. **c** Recurrence Quantification Analysis for the sleep-wake states in **a** and **b** panels using 5 RQA metrics (titles in panels). For each metric, box-plots are constructed from the results of 12 animals and 24 sessions (outliers are not shown).* = *P* < 0.001, ** = *P* < 0.0001, * * * = *P* < 0.00001 (corrected by multiple comparisons).

We find that these results also hold when we divide the frontal-cortex data into specific areas, as well as when we compare them to population activity from the hippocampus (Fig. 3, Table S2); demonstrating the robustness in our conclusions. We analyse data from the secondary motor-cortex (M2), medial pre-frontal cortex (mPFC), orbito-frontal cortex (OFC), and the anterior cingulate cortex (ACC), including data from the CA1 hippocampus region. For these selections, our analyses consistently show that neuronal spiking-activity decreases its temporal complexity during SWS (Fig. 3). In addition, we find no significant differences when we compare the RQA metrics across cortical locations: RR (*P* = 0.68), DET (*P* = 0.39), LAM (*P* = 0.69), TT (*P* = 0.21) and DIV (*P* = 0.46).

**Figure 3:**
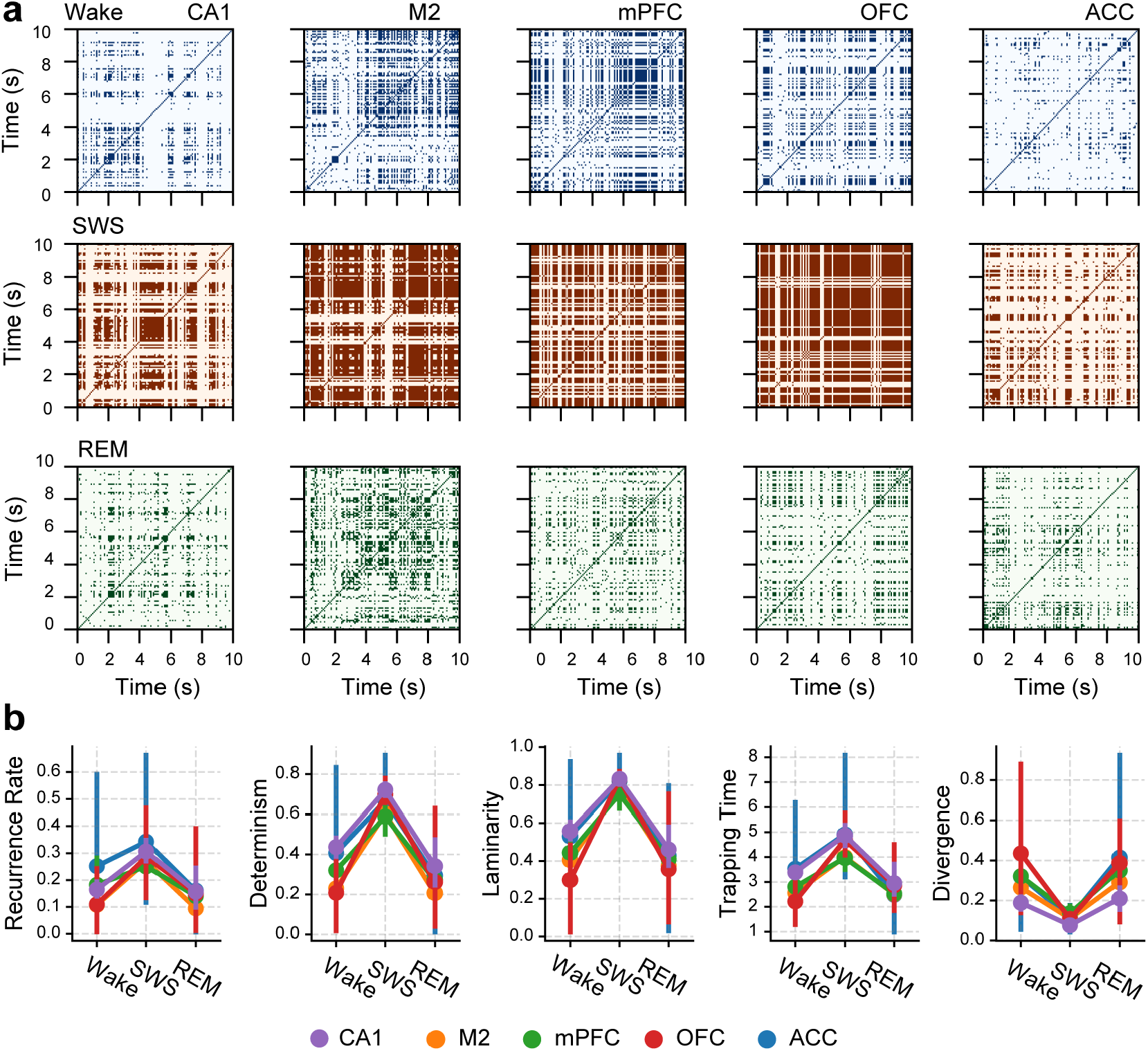
Recurrence analysis for different cortical locations. **a** Examples of recurrence plots for 10 s windows of the binned spike-trains (50 *ms* bins) for different locations during Wake, SWS, and REM sleep. Panels in each column correspond to the Hippocampus (CA1), secondary motor-cortex (M2), medial prefrontal cortex (mPFC), orbito-frontal cortex (OFC), and the anterior cingulate cortex (ACC). **b** RQA metrics for all cortical locations in each sleep-wake state, where filled circles are the population averages and error bars show the corresponding 95% confidence interval.

### OFF-periods explain the complexity changes during SWS in the neocortex

We now study whether the existence of trapped trajectories is correlated to neocortical OFF-periods [26, 28, 29], i.e., ~ 85 *ms*-long periods where almost all neurons remain silent [26]. EEGs reflect coherent activity of pyramidal neocortical neurons [33], where OFF-periods are evidenced in the EEG by the presence of slow waves (0 – 4 *Hz*) [26, 28, 30, 34]. Therefore, a correlation between trapped trajectories and OFF-periods can provide a physiological mechanism for the loss of complexity during sleep.

From Fig. 4**a** and **b**, we can see that OFF-periods (red curve) match the times when recurrent trajectories are in a trapped state (black curve). In general, we find a significant correlation between the time-series of OFF-periods and that of the trapped recurrences: *R* = 0.766 ± 0.02, with *P* = 0 (lower than the 64-bit computer’s float-point) for all sessions. This means that ~ 75% of the SWS trapped-recurrences capture OFF-period dynamics, while the remaining ~ 25% is comparable to the Wake/REM recurrences.

**Figure 4:**
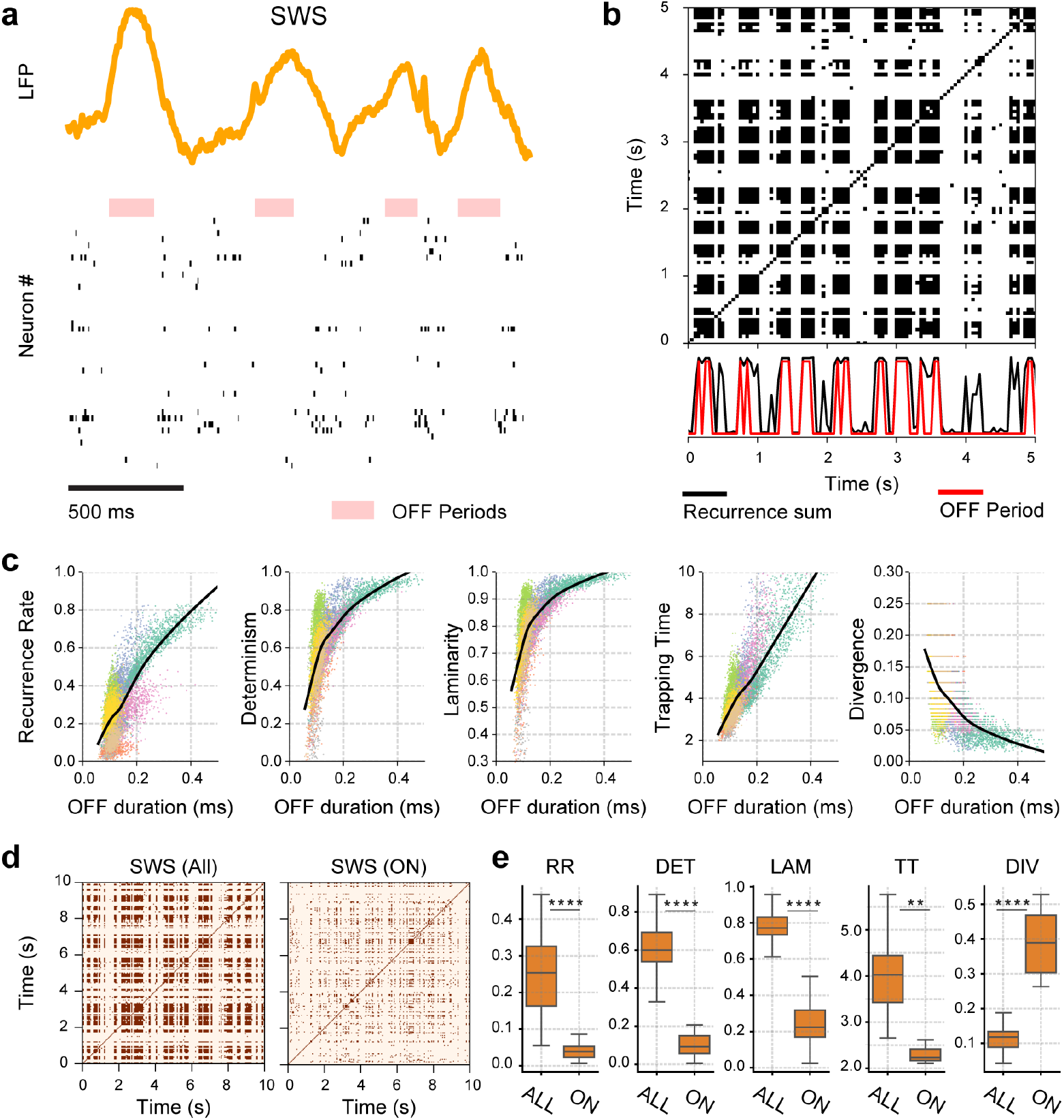
Correlation between recurrent spiking activity and OFF-periods in the neocortex during SWS. **a** Example of a local field potential and spiking activity for a representative animal, where OFF-periods are highlighted by shaded areas. **b** Resultant recurrence plot after binning 10 *s* of spike-trains in 50 *ms* windows, where the number of recurrences in time (sum over columns) is shown in the bottom panel together with the OFF-periods (shaded areas from panel **a**). **c** Correlation between RQA metrics – as those in Fig. 3 – and OFF-period’s average duration; solid lines indicate the LOWESS regression estimate and colours indicate different sessions. **d** SWS recurrence plot containing OFF and ON periods (All), and discarding the OFF-periods (ON) from the spike-trains. **e** Recurrence Quantification Analysis using 5 RQA metrics (titles in panels).** = *P* < 0.001, * * ** = *P* < 0.00001

Moreover, we find that RQA metrics correlate with the mean duration of OFF-periods (Fig. 4**c**). The reason for this is that, the longer the system spends in a silent state, the more it stays trapped in a recurrent state, translating to square-like patterns in the recurrent plots. This implies that RR, DET, LAM, and TT are positively correlated with the OFF-period’s average duration (Fig. 4**c**). Conversely, because of the trapped recurrences, the unpredictability of the system decreases, giving a negatively correlated DIV with the OFF-period’s average duration (Fig. 4**c**). Overall, *P* < 1 × 10^−2^ for the LOWESS regression in any of the recording sessions. In support of these results, we find that if we calculate RQA metrics excluding the SWS OFF-periods (i.e., employing only ON-periods), the complexity reduction is lost (Fig. 4**d,e**).

### OFF-periods account for the loss of complexity in field recordings

In order to see how the RQA results from the population activity translate to field recordings, we create synthetic local field-potentials (sLFP) to compare with real LFP recordings (Fig. 5). We construct a sLFP from the spiking activity of excitatory neurons (LFPs primarily reflect dendritic excitatory post-synaptic potentials [33]), assuming that each spike generates an exponentially decreasing post-synaptic potential (PSP) (Fig. 5**a**) and that the field activity arises from the average PSPs. Namely, we average the PSPs over the population of neurons at each time in order to obtain the instantaneous sLFP. We find that sLFPs have asynchronous low-amplitude activity during Wake and REM sleep, but have synchronous activity during SWS with periodic high-amplitude waves (Fig. 5**b**). These waves correspond to slow-oscillations (0-4 *Hz*, Fig. 5**c** top), coherent to the LFP activity (Fig. 5**c** bottom).

**Figure 5:**
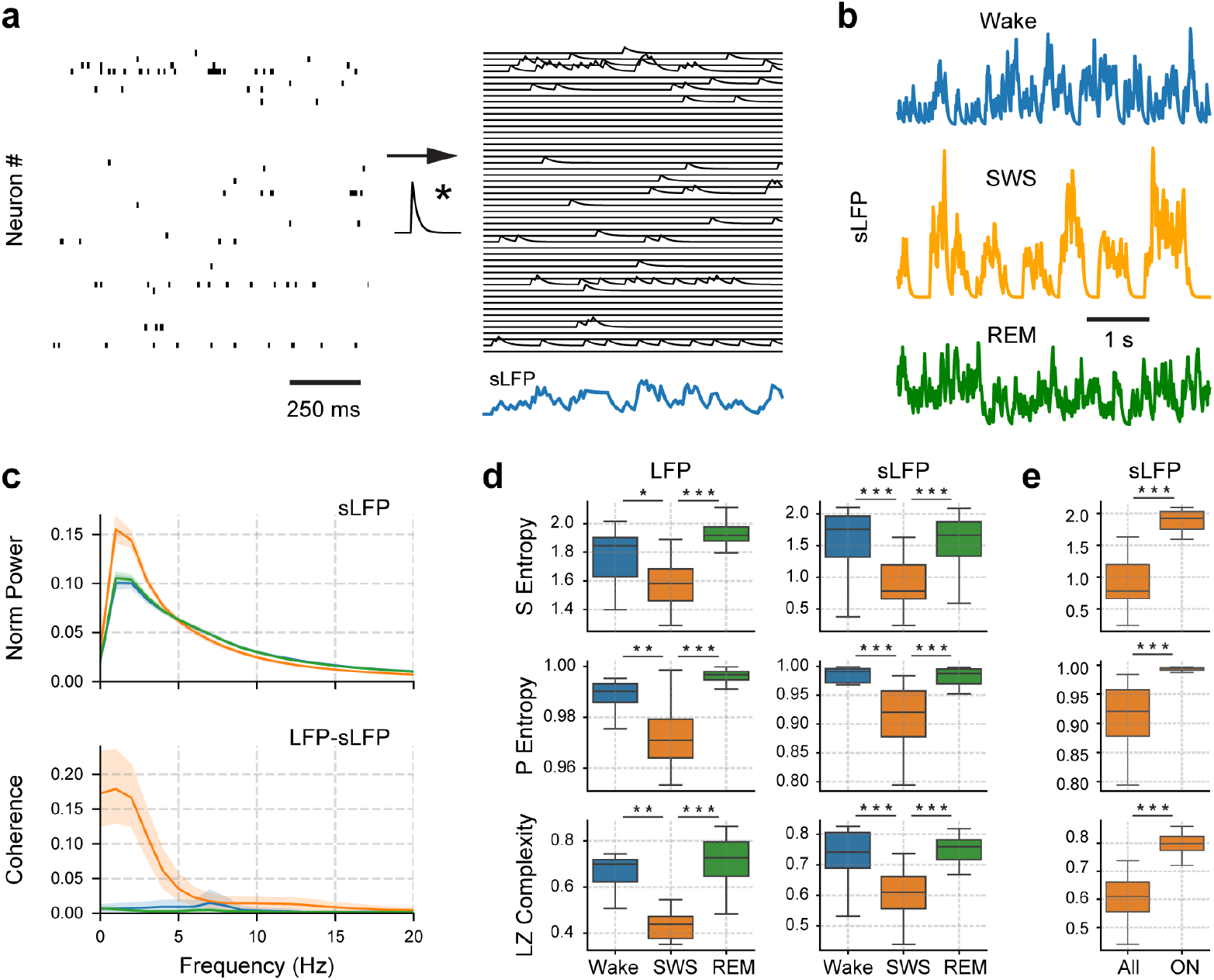
Construction and analysis of synthetic local field potentials (sLFP) during the states of Wake, SWS, and REM sleep. **a** Example of the sLFP creation. A convolution between the neurons’ raster plot and a decreasing exponential function gives a continuous representation of each neuron’s activity. The sLFP (bottom signal) is the ensemble average of these signals. **b** Examples of sLFP resulting from Wake, SWS, and REM sleep population activity. **c** sLFP power spectra (top panel) and coherence between sLFP and LFP (bottom panel) for the different sleep-wake states (colour code). **d** Sample Entropy (top), Permutation Entropy (middle), and Lempel-Ziv Complexity (bottom) of the LFPs and sLFPs in each state for all sessions and animals. **e** sLFP entropies for the SWS signals (All) and the SWS containing only ON-periods (ON), i.e., without OFF-periods. * = *P* < 0.05, ** = *P* < 0.01, * * * = *P* < 0.001

We quantify the temporal-complexity of LFP and sLFP by using Sample Entropy (SE), Permutation Entropy (PE) and Lempel-Ziv Complexity (LZ) [4–11, 14, 15, 18, 32]. Results from this quantification are shown in Fig. 5**d**, which are consistent for all metrics. Firstly, we confirm that the LFP activity is significantly less complex during SWS than during REM or Wake (left panels in Fig 5**d**; Table S3). These results are consistent with previously reported results for EEG and ECoG data [4–13]. Secondly, we obtain similar temporal-complexity values for the sLFP (right panels of Fig. 5**d**; Table S4), which also show significant decrease during SWS. Finally, we test whether OFF-periods are necessary for the complexity reduction during SWS. We construct sLFP for SWS only employing ON-periods; that is, excluding OFF-periods. Panel **e** in Fig. 5 shows that when we re-analyse the ON-period sLFP, the decrease in complexity during SWS is lost (Table S4). In fact, the ON-period SWS has significantly higher levels of complexity than the SWS sLFP containing OFF-periods; values that are comparable to those from Wake or REM states. Therefore, we conclude that OFF-periods are necessary for the complexity reduction observed in field recordings.

### Universal avalanches govern the spiking periods across the sleep-wake states

Our results from Figs. 2, 4, and 5 show that the temporal complexity of the cortex decreases during SWS due to the presence of OFF-periods. Now, we complement those results by analysing spike avalanches in the frontal cortex occurring only during ON-periods, which, in principle, can contribute to decrease the complexity of SWS.

Avalanches are cascades of activity in quiescent systems [23, 35, 36, 44, 45]. In our case, active spiking periods within a brain region; hence, by definition spike avalanches exclude OFF-periods. An avalanche starts in a time-bin that contains at least one spike, happening after a time-bin without spikes, and lasting until another time-bin without spikes is reached. For example, Fig. 6**a** shows a neuronal population exhibiting an avalanche, where the time-bin is defined by the average inter-spike interval (ISI). Two parameters commonly characterise an avalanche: its size, i.e., total number of spikes, and its duration, i.e., time from start to finish. The avalanche statistics for each sleep-wake state are derived from the probability distribution of these parameters [35, 36].

**Figure 6:**
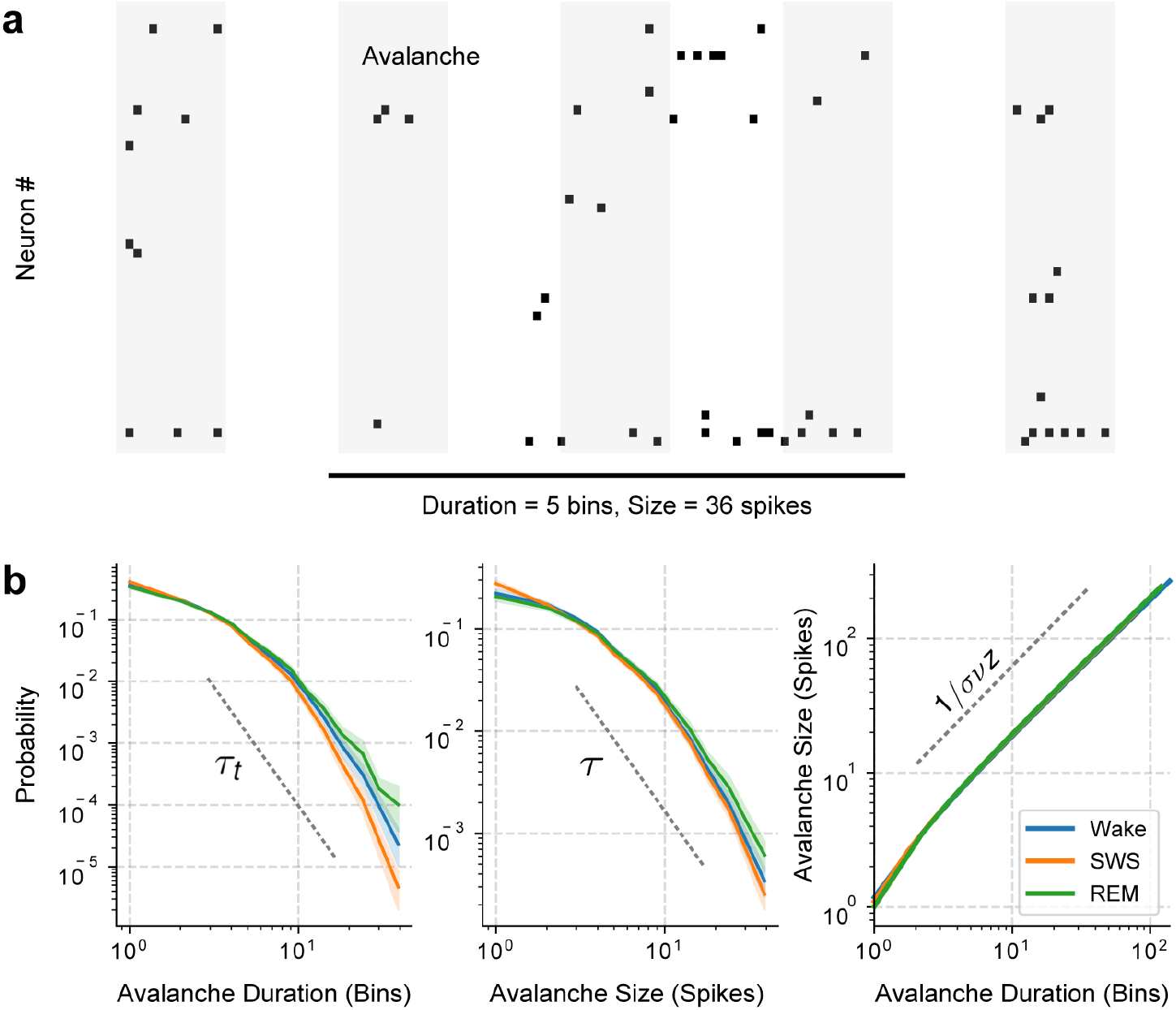
Avalanche distributions for the states of Wake, SWS and REM sleep. **a** Example of how to compute a neural avalanche from a raster plot. The average inter-spike interval (ISI) is used to bin (shaded areas) the raster plot and count the number of spikes per bin. **b** Left: distribution of avalanche durations, where we estimate a *τ_t_* exponent. Middle: distribution of avalanche sizes, where we estimate a *τ* exponent. Right: avalanche size as a function of its duration, where we estimate the 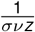 exponent. For each state, mean distributions are shown in solid lines with a shaded area signaling the 95% confidence interval.

Our findings show minimal differences between the probability distribution of avalanches’ duration or size during Wake, REM sleep, or SWS (left and middle panels of Fig. 6**b**). More importantly, these distributions collapse to the same scaling function (right panel of Fig. 6**b**). This implies that the spiking periods have a universal behaviour, independently of the sleep-wake state. In particular, the scaling function for the neuronal spiking avalanches approximates a power-law behaviour, which can be characterised by its exponent, 1 /*σνz*. We find that 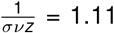 for all sleep-wake states (inter-quartile range, *IQR* = 0.05) with *P* = 0.21, implying that differences between sleep-wake states are not significant. Similarly, the power-law exponents for the avalanche duration (*τ_t_*) and size (*τ_t_*) are related by 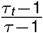 which for the neocortical neurons and all sleep-wake states is 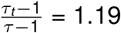 (*IQR* = 0.33, with *P* = 0.31).

We also calculate the branching parameter for the avalanches. This parameter quantifies how spikes propagate during an avalanche, measuring the average of the number of spikes at time *t* + 1 given a single spike at time *t*, i.e., 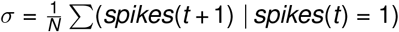 [23]. The branching parameter (*σ*) yields a median of *σ* = 1.01 ± 0.01, without differences across states (*P* = 0.11). Same as with the avalanche statistics during SWS, we calculate the branching parameter excluding OFF-periods.

These results show that complexity differences in the sleep-wake states originate from OFF-period’s distribution. In fact, once the spiking activity is initiated, it follows an avalanche behaviour with a universal scaling relationship, irrespective of the sleep-wake state. These results restrict the possible mathematical models which can describe cortical-dynamics, since the model must be able to reproduce OFF-periods (during SWS) and the universal avalanches appearing for any state.

### Critical branching model for the spiking activity in the cortex

Here, we show how our results can be captured by a critical branching model [37], known to exhibit universal behaviour with similar exponents to our *in-vivo* results. The critical branching model consists of interacting discrete units that evolve in time, whose internal state may be resting, spiking, or refractory; thus modelling the neuron’s basic states. The branching parameter, *σ*, controls the probability of a spike from unit A at time t affecting unit B at time *t* +1. When *σ* = 1, the system is critical, having a phase transition from a sub-critical quiescent state for *σ* < 1 (activity dies out after a small transient) to a super-critical active state for *σ* > 1 (activity is self-sustained). The units evolve according to the excitation coming from neighbouring units as well as due to a noisy component (set by a Poisson distribution), which can randomly change the state of any unit at any given time. The interplay between units interacting due to branching and the noisy substrate recreates a network of higher-order neurons that receives inputs from lower areas. To reproduce a SWS state, we add to the branching model a periodic silencing of the noise for some (adjustable percentage of) units in order to model OFF-periods.

Figure 7**a** shows an example of the resultant spike trains for the branching model (left panel) and our modified version (right panel); that includes a periodic silencing of the Poisson noise. These results are obtained using 50 units (similar size to the experimental ensembles recorded) and setting the branching parameter at the critical point: *σ* = 1, in agreement with the experimental value measured. Their respective recurrence plots are shown in Fig. 7**b**. On the modified branching model, we periodically silence the noise input arriving to a given set of units during a 250 *ms* interval (similar to Ref. [35]), and call it as Critical + OFF-periods. This external forcing is enough to drive the field activity to a synchronised state of inactivity (Fig. 7**a**), trapping the population activity’s trajectory into recurrent square-like patterns (Fig. 7**b**), similar to the experimental results from the neocortex and hippocampus (see Fig. 2**b** and 3**a**).

**Figure 7:**
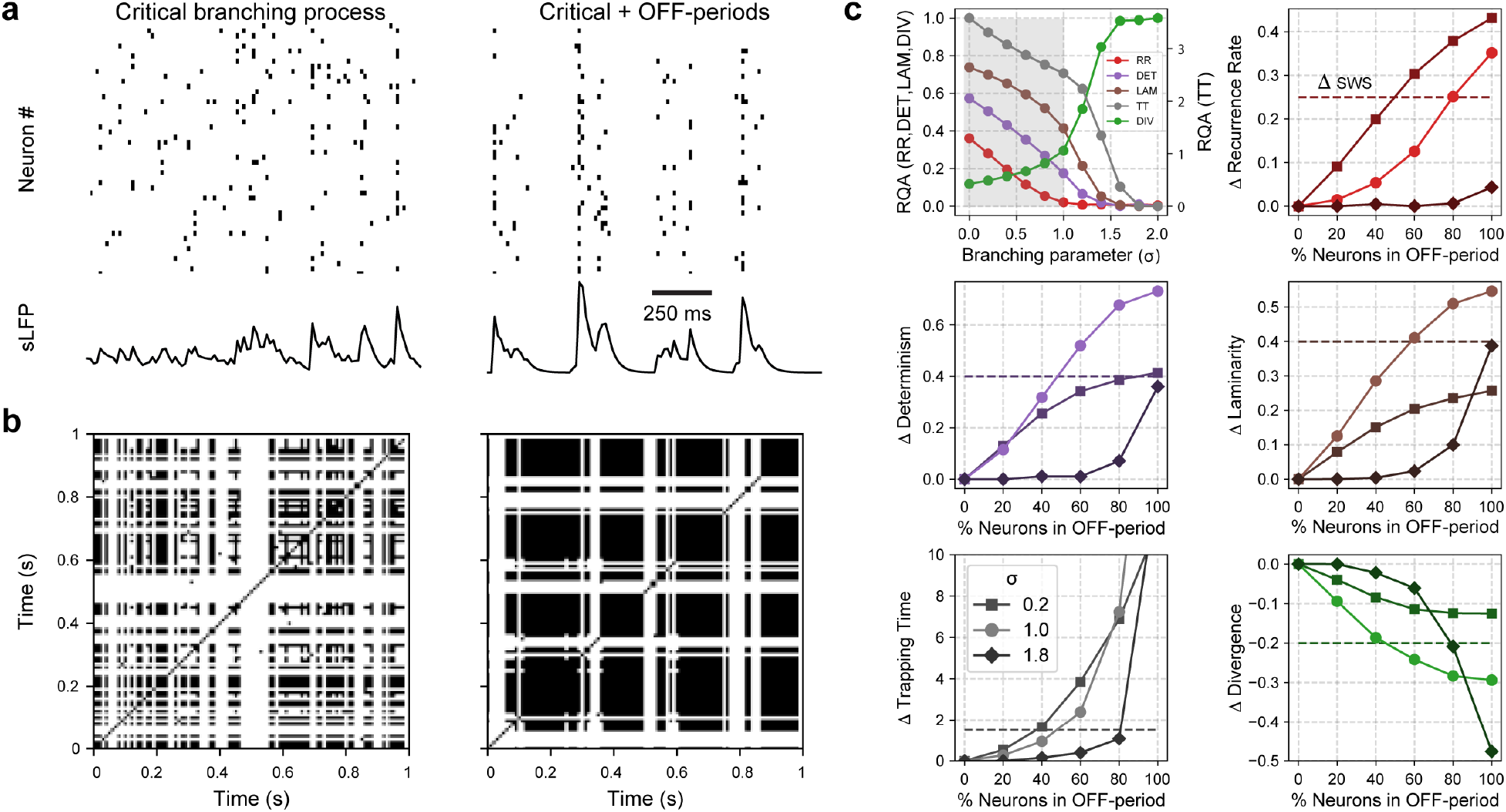
Modified critical branching model reproduces neuronal activity of Wake and SWS. **a** Population activity (raster plot) obtained from the model for 50 neurons and their synthetically generated local-field Potential (sLFP) (see Fig. 5). Left: model without noise silencing and tuned to its critical point, where the branching parameter *σ* = 1. Right: model tuned to *σ* = 1 with an external periodic-silencing of the noise. **b** Resultant recurrence plots for the data from panels **a**. **c** RQA metrics as function of *σ* and the percentage of neurons with the noise periodically silenced. Top left: RQA metrics for the (pristine) critical branching model as function of branching parameter *σ* (see Methods). Grey [white] band shows the subcritical [super-critical] phase. Remaining panels: differences between RQA metrics for the model with and without periodic-silencing of the noise, Δ, as a function of the percentage of neurons having their noise silenced. Colours for Δ RQA metrics follow those from top left panel **c**. The horizontal dashed line shows the Wake-SWS difference for the experimental results. Note that in all cases, circles show the critical model

We use RQA to quantify the differences between the branching model and our modified model, which has a periodic noise-silencing. Results are shown in Fig. 7**c**. The top left panel contains RQA metrics for the branching model with 50 units as a function of *σ*, where the shaded area signals the sub-critical phase. For *σ* = 1, the model has RR≃ 0.02, DET≃ 0.2, LAM≃ 0.4, DIV≃ 0.3, and TT≃ 2.5, which are comparable to the average RQA values of Figs. 2**c** and 3**b** during Wake and REM sleep. The remaining panels show the change in the RQA metrics when the periodic noise-silencing is added to the model – changes are shown as a function of the percentage of units having their noise periodically silenced. By a horizontal dashed line, we also include the SWS RQA metric relative to the critical branching model value. In other words, this relative SWS metric is found taking the value obtained from the experimental average RQA metric shown in Fig. 3**b** and subtracting the critical branching model RQA metric from the top left panel in Fig. 7**c**. Using this, we can find the percentage of units with periodically-silenced noise that are needed to reproduce the experimental values found for SWS.

We find that as the number of units with an OFF-period increases, RQA metrics cross those observed during SWS from the *in-vivo* recordings (horizontal dashed-line). When the model is at the critical point, *σ* = 1, it is enough to apply a periodic noise-silencing to 40 – 60% of the units to reproduce the RQA values during SWS (intersection of the Δ RQA metrics with the corresponding horizontal dashed-lines); with the exception of RR, which requires 80%. On the contrary, both the sub- and super-critical models need a considerably larger percentage of silencing to reproduce the observed SWS values – between 80 −100%. Therefore, these results imply that: i) the branching model needs to be close to *σ* = 1 (criticality) to reproduce the recurrent properties observed during Wake or REM sleep, and ii) that the inclusion of a periodic noise-silencing to 40 – 60% of the units reproduces the recurrent properties observed during SWS.

## Discussion

It has been widely reported that during SWS, the temporal complexity of field recordings decreases [4–12]. However, the reason behind this reduction remained elusive. Here, we show: i) that the presence of OFF-periods in neuronal population activity correlates with the complexity reduction of the LFPs during SWS (Fig. 4); ii) that the existence of OFF-periods is necessary for this complexity reduction (Figs. 4, 5 and 6); and iii) that introducing OFF-periods to a critical branching model is sufficient (enough) to reproduce the SWS characteristics from the *in-vivo* recordings (Fig. 7).

### Recurrence quantification analysis to study cortical population dynamics

We analyse the spiking activity from *in-vivo* population recordings during the sleep-wake states, both in the neocortex and hippocampus (Figs. 2 and 3). We employ RQA to retrieve the main dynamical features of the population activity from a 2*D* recurrence plot and quantify its complexity (Fig. 1). A significant advantage of RQA over methods that require dimensionality-reduction, is that it is robust to parameter tuning (e.g changing the recurrence tolerance or the spike count bin-width, Fig. S1), is computationally efficient (10 s windows are enough to find differences between states), and it keeps results and interpretations clear.

RQA also has parallelisms to topological data-analysis, such as persistent homology, which relies on studying the topology of high-dimensional cloud of points (manifold) corresponding to the system’s evolution in phase-space [19, 38]. For example, the anterior nucleus of the thalamus shows a ring-like structure in phase space during Wake and REM sleep, but not during SWS [38]. However, we find that the neocortical phase-space attractor is irreducible to a low-dimensional structure and remains unchanged throughout the sleep-wake cycle (Fig. S2).

### OFF-periods reduce the complexity of cortical activity during SWS

We find that the evolution of the population spiking activity is significantly altered during SWS, in contrast to the unchanged attractor’s topology. In particular, we show that the alteration of the cortex’s dynamics during SWS is due to the presence of population OFF-periods (Fig. 4), allowing us to explain two observations.

On the one hand, slow waves (0.1 – 1 *Hz*) and delta waves (1 – 4 *Hz*) have been associated to the loss of complex neuronal-interactions during sleep [24, 31, 32]. This observation is consistent with our power spectrum and coherence results (see Fig. 5**C**), which further confirms the relationship between OFF-periods and slow waves [26, 30, 34, 39]. Moreover, it was speculated that the nature of the undergoing oscillation could constrain the firing pattern diversity [40]. We confirm this by showing that the oscillation’s neural substrate determines the overall complexity.

On the other hand, it had been shown for individual neurons that the complexity of firing patterns decreases during SWS [6]. This decrease can be explained by OFF-periods, as their appearance causes neurons to remain silent during synchronous intervals, making their overall firing-patterns less complex. However, we note that when analysing the firing patterns of our neurons independently, we find that a considerable number of neurons maintain a complex pattern even during SWS (see Fig. S3). Therefore, we argue that the complexity reduction is a population-level phenomena, which we support by finding the complexity differences between Wake/REM and SWS increase as the number of simultaneously recorded neurons included in the analysis grows (see Fig. S4).

We highlight that we also observe a complexity reduction during SWS in the hippocampus, in spite of being an area whose OFF-periods differ from the ones described in the neocortex [30, 41]. During SWS, hippocampal neurons oscillate between long quiescent stable periods (without clear membrane hiperpolarization), and bursts of spiking activity (during sharp-wave ripples). Contrarily, neocortical neurons oscillate between stable periods of spiking activity and unstable periods of quiescense (associated with hiperpolarization). In spite the difference in the underlying cellular mechanisms, both population activities are consistent with excitable dynamics [41]. Therefore, in both areas, the transition to an excitable population regime during SWS leads to recurrent patterns, which cause the complexity reduction.

### Measuring complexity from field recordings

The complex nature of brain activity, and its reduction during unconscious’s states, has been reported using classical neuroscience approaches [42, 43]. For instance, the EEG power spectrum shows a power-law decay, *f^−α^*, for a broad frequency range *f*, referred as 1/*f* noise. Interestinlgy, this exponent becomes smaller than 1 (i.e., a more pronounced decay) during sleep and anaesthesia [42, 43]. We also find this in LFPs and ECoGs (Fig. S5), where we get similar decay exponents during SWS and REM (*α_sleep_* ≃ 2) but different from Wake’s exponent. However, the exponent differences is significantly higher in ECoGs, where *α_wake_* ≃ 1, than in LFPs, where *α_wake_* ≃ 2 (as during sleep). This could point to the presence of extra-neural sources during Wake that alter the ECoG’s power-spectrum decay, but disappear at the LFP recording level.

These observations are also consistent with our RQA of the rat’s cortex neuronal activity. We justify that OFF-periods are responsible for reducing the complexity of field recordings during SWS by generating synthetic field recordings from the population activity (Fig. 5). From this synthetically constructed LFP, we see that complexity reductions during SWS are lost when we eliminate the OFF-periods from the construction (see Fig. 5**e**). Because OFF-periods and slow-oscillations promote the appearance of low frequencies, it is expected that the power-spectrum decay becomes steeper (i.e., *α* grows).

### Spiking periods show universal dynamics across states

An important result is that our analysis shows that population activity during SWS’s ON-periods is indistinguishable to the activity during Wake or REM. This strengthens our claim that OFF-periods are responsible for the change in SWS complexity. As our results from RQA (Fig. 4), field complexity metrics (Fig. 5), and avalanche statistics (Fig. 6) show, while spiking activity is occurring (i.e., during ON-periods), SWS behaves exactly as (if not more complex than) Wake or REM.

Specifically, we show that neuronal avalanches of length *t* contain an average of *f*(*t*) spikes, where *f* is a scaling function independent of the sleep-wake state. This means that avalanches from the frontal cortex of rats, follow a universal behaviour, previously reported in the visual cortex [35]. We find that the scaling exponent 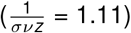 and the exponent relating the avalanche statistics 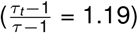 are relatively similar, which is an expected relationship when a system is close to criticality [36]. Therefore our results support the hypothesis that complex cortical activity arises from near-critical dynamics [23, 35, 36, 44–49].

### OFF-periods are sufficient to reproduce the complexity reduction in a critical model of the cortex

To complement our *in-vivo* results, we show that the introduction of OFF-periods into a critical model is sufficient to generate a SWS-like complexity decrease. We achieve this by periodically silencing the noise that is inherent to the model. This periodic silencing reproduces OFF-periods, which are generated primarily from pre-synaptic inhibition of excitatory inputs to principal cells [50]. In fact, a similar strategy was employed to model anaesthesia [35], suggesting that similar dynamics might underlie the complexity reduction during that state.

We find that near the critical point of the model, the percentage of neurons we need to force to get a SWS-like state is optimal with respect to either the sub- or super-critical state. For instance, silencing the input to 40 – 60%, creates a complexity reduction similar to that observed experimentally. This further adds to the idea of criticality in the brain, which would explain the increased complexity [22], information processing and transmission [23], and dynamical range [46].

## Methods

### Datasets

We analyse 3 datasets: Watson et al. (population activity neocortex, available at http://crcns.org/data-sets/fcx/fcx-1) [51], Grosmark and Buzsaki (population activity hippocampus, available at http://crcns.org/data-sets/hc/hc-11) [52], and Gonzalez et al 2019 [9] (ECoG neocortex, available at request). The reader is referred to the original publications for details about experimental methods. We provide a summary below.

For the frontal cortex dataset, silicon probes were implanted in frontal cortical areas of 11 male Long Evans rats. Recording sites included medial prefrontal cortex (mPFC), anterior cingulate cortex (ACC), pre-motor cortex/M2, and orbitofrontal cortex (OFC). Recordings took place during light hours in the home cage (25 sessions, mean duration of 4.8 hs ± 2.2 std). We note that we exclude *BWRat19_032413* from the analysis, as it did not contain REM sleep. Data was sampled at 20 *kHz*. To extract LFPs, recordings were low-passed and resampled at 1250 Hz. To extract spikes, data was high-pass filtered at 800 *Hz*, and then threshold crossings were detected. Spike sorting was accomplished by means of the KlustaKwik software. Sleep-wake states were identified by means of principal component analysis. SWS exhibited high LFP PC1 (power in the low < 20 *Hz*) and low EMG. REM sleep showed high theta/low EMG cluster, and a diffuse cluster of low broadband LFP with higher EMG. Wake showed a diffuse cluster of low broadband LFP, with higher EMG, and a range of theta. OFF periods were extracted as periods of population silence lasting at least 50 *ms* and no more than 1250 *ms*. Conversely, ON periods consisted of periods of population firing between OFF periods with at least 10 total spikes and lasting 200 – 4000 *ms*. For more details, see [51].

For the hippocampus dataset, silicon probes were implanted in the dorsal hippocampus (CA1) of 4 male Long Evans rats (7 recordings total). LFP and spikes were extracted similarly to the cortical dataset. A similar criterion as before was employed to identify the sleep-wake states; for a full description, see [53].

For the ECoG dataset, 12 animals with 7 steel screw electrodes placed intracranially above the dura matter were analysed. The recorded areas included motor, somatosensory, visual cortices (bilateral), the right olfactory bulb, and cerebellum, which was the reference electrode. Data was sampled at 1024 *Hz*, employing a 16 bits AD converted. The states of sleep and wakefulness were determined in 10 s epochs. Wake was defined as low voltage fast waves in the motor cortex, a strong theta rhythm (4–7 *Hz*) in the visual cortices, and relatively high EMG activity. NREM sleep was determined by the presence of high voltage slow cortical waves together with sleep spindles in motor, somatosensory, and visual cortices associated with a reduced EMG amplitude; and REM sleep as low voltage fast frontal waves, a regular theta rhythm in the visual cortex, and a silent EMG except for occasional twitches. Additional visual scoring was performed to discard artefacts and transitional states.

### Recurrence quantification analysis

Prior to analysing the recurrences [54–56], we bin the spike data to 50-ms firing count bins.

Given a trajectory 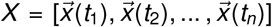 composed of the distinct time measurements of the vector 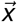, whose entries are the different firing counts of each neuron, a recurrence plot is defined by a matrix *R* whose entries are:

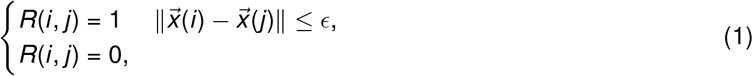

where *ϵ* > 0 is the tolerance defining closeness. Thus, a recurrence between two time points would only occur if the system was located in a similar region (state) of the phase space at the respective times (up to an error of *ϵ*). In our case, *ϵ* is set to 1 standard deviation of the population firing counts.

To quantify the patterns arising from recurrence plots, we employ common measures form recurrence quantification analysis (RQA). The metrics (defined below) are: recurrence rate (RR), determinism (DET), laminarity (LAM), trapping time (TT) and divergence (DIV)

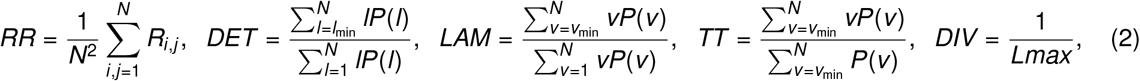

where *P*(*l*)[*P*(*v*)] indicate the probability of finding a diagonal [vertical] line of length *l*[*v*], and *Lmax* indicates the longest diagonal line, excluding the identity line.

### Synthetic LFPs and field complexity measures

To construct sLFPs, we convolve the spike times (125 Hz sampling bin) of each excitatory neuron *S_n_* by an exponentially decreasing kernel, to obtain the convolved spikes *C_n_* of the nth neuron.

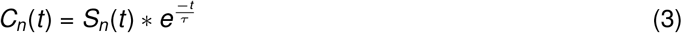

the symbol * is the convolution operator and *τ* is the exponential time constant. *τ* is set to 24 ms for all putative excitatory neurons, which is a reasonable mEPSP time-constant for a pyramidal neuron in the frontal cortex [58]. This selection of *τ* is also validated internally, as it allows to recover important LFP features in the sLFP.

We obtain the sLFP is averaging the convolved spikes acoss neurons.

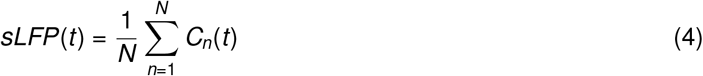

*N* is the total number of simultaneously recorded neurons.

We analyse the sLFPs power spectrum by means of Welch’s algorithm employing the *signal.welch* scipy python 3 function (https://www.scipy.org), setting a 1s moving Hanning window, no overlap, and a 1 Hz frequency resolution. To get the sLFP-LFP coherence, we employ the *signal.coherence* scipy function. Same parameters as the frequency spectrum. We average the LFP across channels and downsample it to 125 Hz prior to estimating the coherence to the sLFP.

To quantify the sLFP and LFP temporal complexity, we employ Permutation Entropy, Sample Entropy and Lempel-Ziv Complexity, *antroPy* python 3 package (https://github.com/raphaelvallat/antropy). Permutation Entropy [57] consists in encoding a time-series {*x*(*t*), *t* = 1, …, *T*}, by dividing it into ⌊(*T* – *D*)/*D*⌋ non-overlapping vectors, *D* is the vector length, which is much shorter than the time-series length (*D* ≪ *T*). Then, each vector is classified according to the relative magnitude of its *D* elements. We employ *D* = 3, and *τ* = 5 (*τ* being the distance between succesive time-stamps in each vector). We then compute the Shannon entropy (SE) [59] to obtain Permutation Entropy. 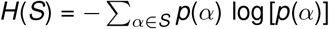, where *p*(*α*) is the probability of finding symbol *α* in the signal (among the set of symbols *S*) and the summation is carried over all possible symbols.

Similar to Permutation Entropy, Sample entropy consists of dividing a time-series {*x*(*t*), *t* = 1, …, *T*} into a series of *D* sized vectors (*X*(*i*)). The Chebyshev distance (*d*) is applied to vector pairs (with different indexes, like *X*(*i*), *X*(*j*)). *A* is obtained as *d*[*X*_*m*+1_(*i*), *X*_*m*+1_(*j*)] < *r* and *B* as *d*[*X_m_*(*i*), *X_m_*(*j*)] < *r*. Sample entropy is defined as: 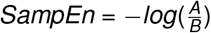 In our case *D* = 3, and we downsample the signals by a factor of 5, in order to match the permutation entropy *τ*.

Lastly, we employ the Lempel-Ziv complexity algorithm. Prior to apply this function, we binarize the sLFP signals by its mean value (all time-points above the signal’s mean are converted to a 1 and all below to a 0) and divided it into 10s windows. Then the Lempel-Ziv complexity is estimated (LZ-76 algorithm), counting the the number of different substrings encountered as the sequence is viewed from beginning to the end. We normalise it as 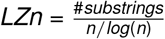, making the metric less dependent on the signal’s length.

### Neuronal avalanches

We quantify neuronal avalanches following previous studies [35, 36]. First multy-unit activity is binned, employing the average ISI. Then, we measure the time (duration) and number (size) of spikes between one empty bin (0 spikes) to the following empty bin. We use the *powerlaw* (https://pypi.org/project/powerlaw/1.3.4/) python 3 package to construct the probability distributions and obtain their exponents (*τ_t_* and *τ*). We also compute the average number of spikes as a function of the avalanche duration, and obtain the 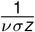 exponent by means of an ordinary least square fit on a log-log scale. We calculate the branching parameter for the avalanches in each state, measuring the average of the number of spikes at time *t* + 1 given a single spike at time *t*, i.e., 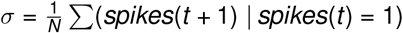 [23].

### Critical branching model

The critical branching model consists of 50 interacting units randomly connected in a Erdos-Renyi topology (each pair of neurons has a 0.01 probability to be connected). Each unit has 3 possible states: resting, firing or refractory. The transition between resting and firing can either occur from the excitation coming from a connected neuron firing in the preceding time, or by the intrinsic Poisson noise that each neuron receives independently. The Poisson noise is set by generating a random matrix whose values come from a [0, 1] uniform distribution, and then setting for each entry a spike if the value is less than 1 – *e*^−λ^ (λ = 0.014). Once a neuron fires, it goes deterministically to the refractory state, in which it cannot be excited again. After 1 step in a refractory state, each neuron goes to the resting state and becomes excitable again. The propagation of spikesis controlled by the branching parameter (*σ*), which regulates the overall excitability of the system. For instance, if neuron *i* fires, the probability that a neighbouring unit fires is defined as 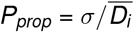, where 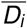 is the average node degree for unit i. To obtain the plots shown in Fig. 7, we employ 1 million iterations from each network, and average the results over 100 trials. To obtain a SWS state, we periodically silenced the Poisson noise coming to the network in a 250ms step with a 4*Hz* frequency.

### Persistent homology

To study the topology of the neural manifolds during the states of sleep and wakefulness, we employ similar procedures as [19, 38]. Briefly, we bin the spike data to 100-ms firing count bins and then reduce its dimensionality by means of the isomap algorithm to a 3d representation. After that, we calculate persistent homology by means of the *ripser* python3 package, limiting the analysis to Betti 0 and 1 numbers. To compare between conditions, we select the most persistent Betti 0 and 1 components for each session in each state.

### Single neuron complexity measures

To quantify the complexity of the firing patterns of single neurons, we employ the Lempel-Ziv complexity of the raster plots (10*s* windows) and the inter-spike intervals (ISIs) entropy. For the ISI entropy, we construct the ISI histogram employing 18 bins equally spaced between 0 and 800 ms. After that, the Shannon entropy is computed as mentioned above. We compute the Lempel-Ziv complexity as defined in the previous section.

### Statistics

We present data as regular boxplots showing the median, the 1st and 3rd quartiles, and the distribution range. Because of the complexity metrics we analyse, we employ non-parametric statistics. We use the Friedman test (available with the *scipy.stats*) to compare the results among states (Wake-SWS-REM) with the Siegel post-hoc test applying the Benjamini-Hochberg false discovery rates correction (available with the *scikitlearn* python 3 package (https://scikit-learn.org/stable/)). We also employ the Wilcoxon sign-ranked test to compare between SWS(All) vs SWS(ON). We set P < 0.05 for a result to be considered significant. In addition to P-values, we also reprt Cohen’s d, which quantifies the magnitude of a result in terms of a standardised difference between conditions, considering an effect size to be large if Cohen’s d is > 0.8. For the power spectrums and avalanche results we present the data as the mean with the 95% confidence interval (obtained through bootstrap sampling). For the correlation analysis, we employ LOWESS regression to fit the best estimate to the scatter plot, by means of the regplot function available at *seaborn* (https://seaborn.pydata.org) python 3 function. As LOWESS regression doesn’t have an associated *P* value, we employ a linear regression for each session and report the result significant only if *P* < 0.05 for all sessions. Additionally, to correlate the OFF-periods to the recurrence sum, we employ the point-biserial correlation *pointbiserialr* available at *scipy* https://scipy.org.

## Code availability

The codes developed to analyse population activity recurrences are freely available at https://github.com/joaqgonzar/Recurrence_population_activity, including the recurrence functions and jupyter notebook examples.

## Acknowledgements

JG acknowledges the support of Comisión Académica de Posgrado (CAP), CSIC Iniciación and PEDECIBA. PT also acknowledges the support of PEDECIBA. ABLT acknowledges the support of CAPES and CNPq.

## Supplementary Material

**Figure S1:**
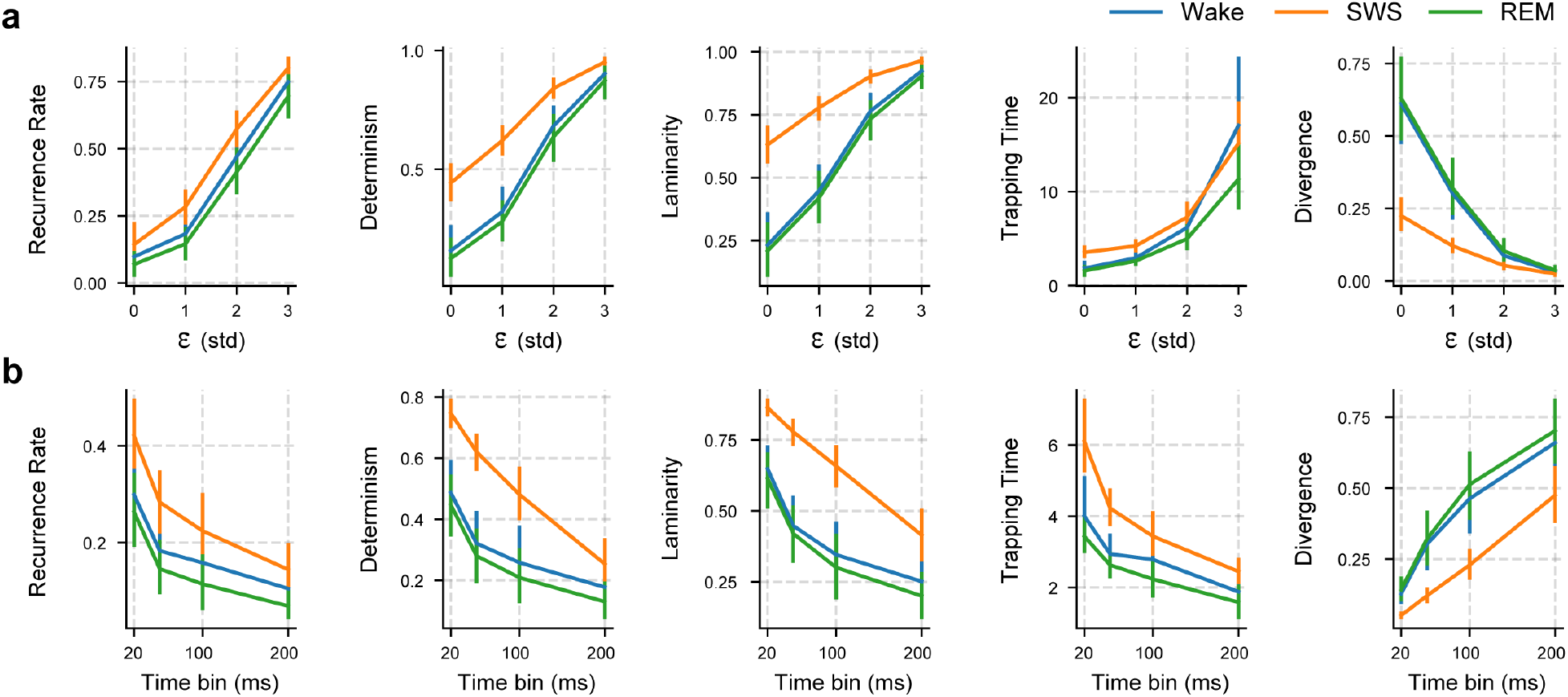
RQA is robust to parameter choice. **a** RQA metrics for different tolerance levels *ϵ* for defining recurrence in phase-space. We vary *ϵ* from 0 std to 4 std of the population firing counts. Setting *ϵ* to 0 means that a recurrence occurs between two times for the exact same neuronal firing pattern. The time bin is kept fix at 50*ms*. **b** RQA metrics for different time binning of the population activity. Time bins are changed from 20 ms to 200 ms in order to define the firing variable for each neuron. The *ϵ* is kept fix at 1 std. The mean and its corresponding 95 % confidence intervals are shown for each plot.

**Table S1:**
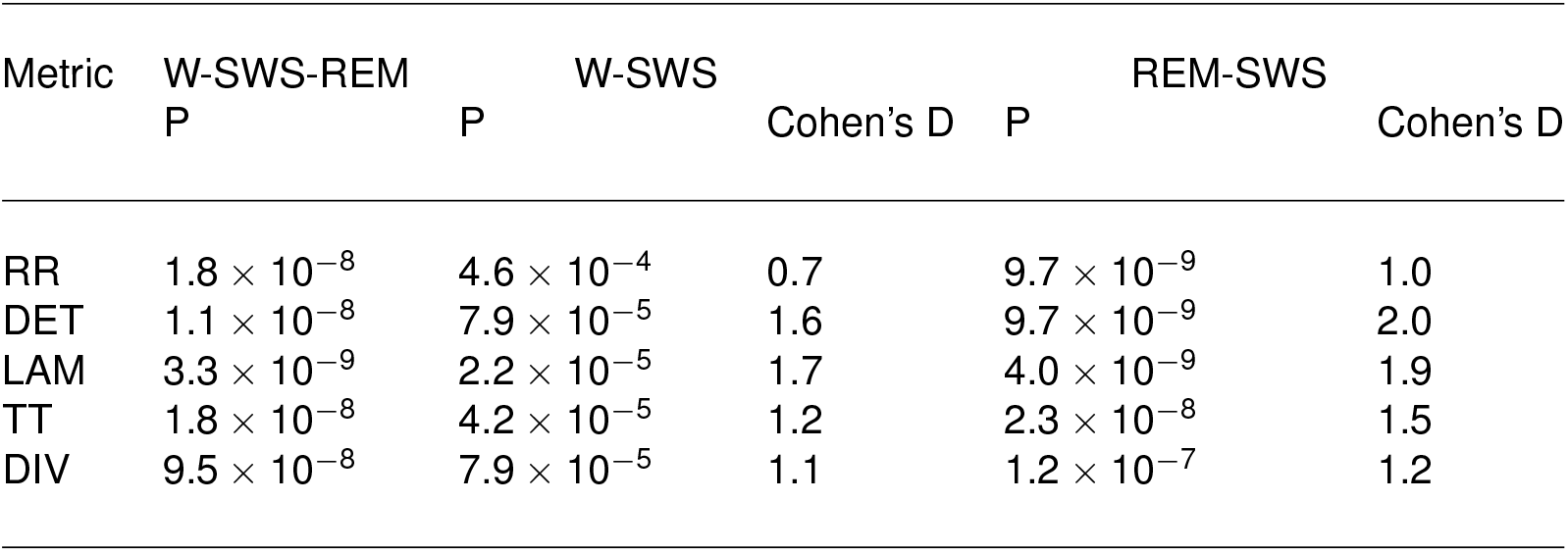
Statistical comparisons for RQA in the neocortex.

**Figure S2:**
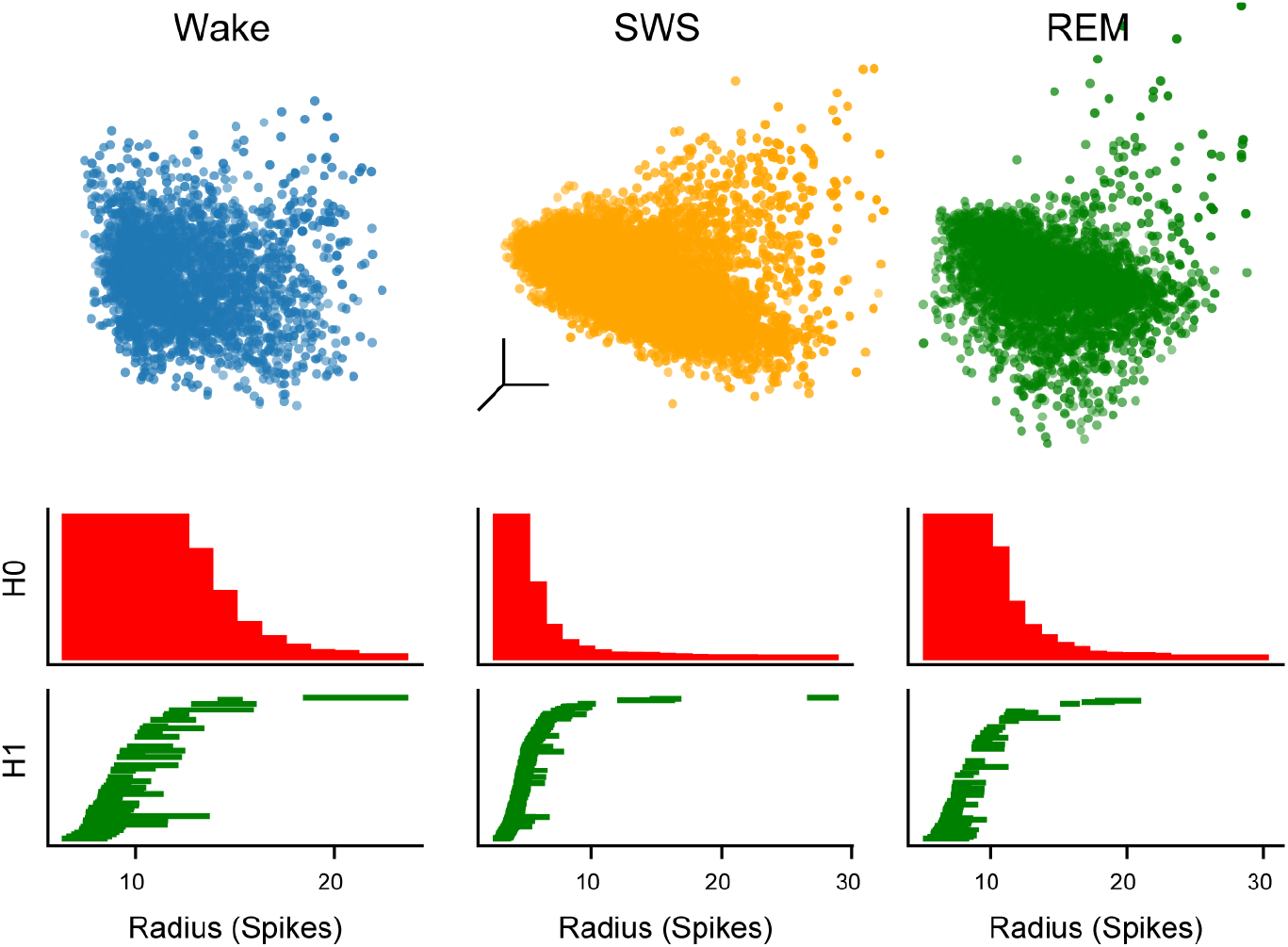
Persistent Homology during the sleep-wake states in the neocortex. Top panels: Point cloud obtained after dimensionality reduction. A representative animal is shown during Wake, SWS and REM sleep. Bottom panels: Betti 0 (HO) and Betti 1 (H1) barcodes for the same animal shown in the top panel. The length of each bar shows the level of persistence of each Betti 0 and 1 component.

**Figure S3:**
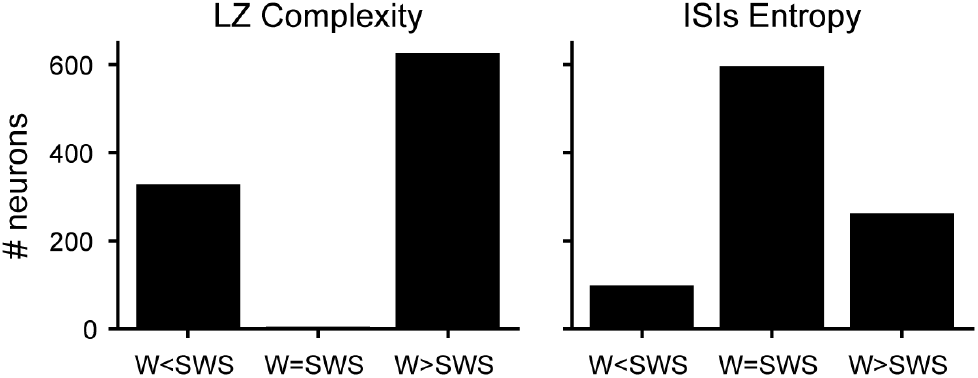
Single neurons deviate from the ensemble behaviour. Complexity of single neuron firing pattern between Wake and SWS. Each bar shows the total number of neurons whose temporal complexity decreased, remained equal or increased. 2 metrics are shown. Left: Lempel-ziv complexity of single neuron binary raster. Right: Inter-spike interval histogram entropy.

**Table S2:**
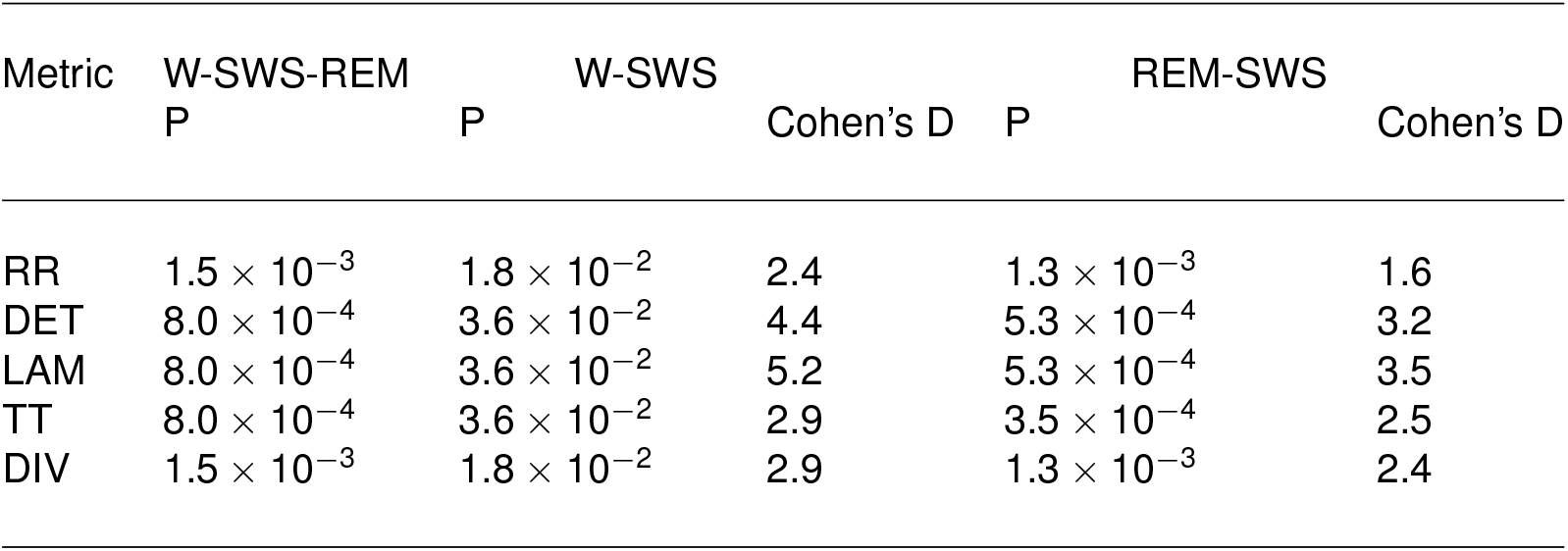
Statistical comparisons for RQA in the hippocampus.

**Figure S4:**
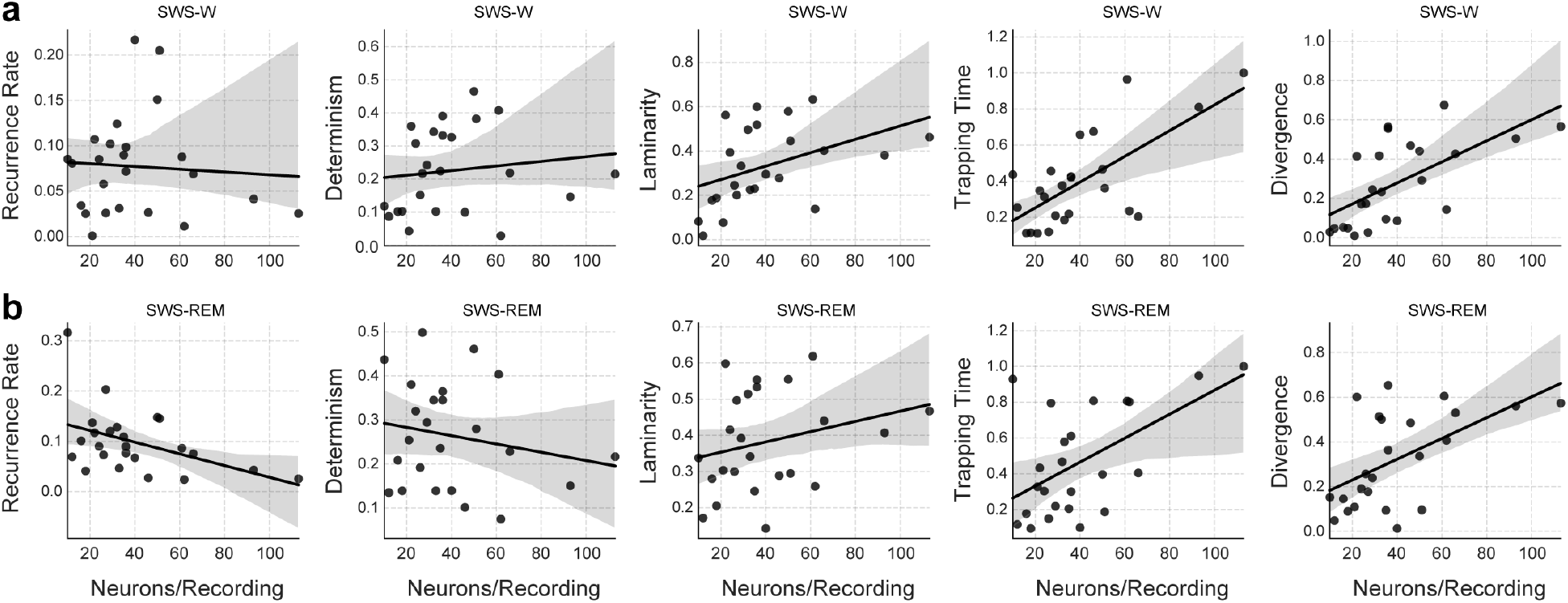
RQA differences between states correlate with the number of neurons recorded. Absolute RQA differences between states as a function of the number of simultaneously recorded neurons. Each dot shows a recording session while the solid line the linear regression estimate with its 95 % confidence interval. **a** shows the SWS-Wake difference, while **b** the SWS-REM difference.

**Figure S5:**
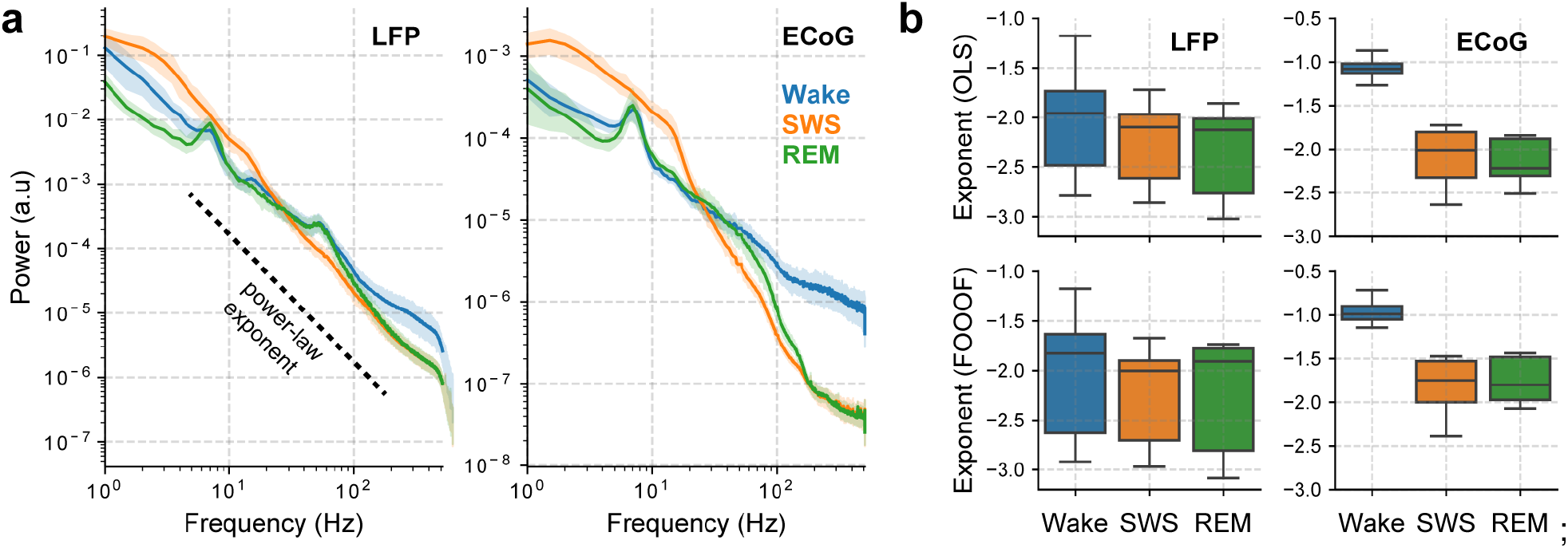
Power spectrum slope differs between states. **a** LFP [ECoG] recordings coming from the frontal cortex [M1 cortex] during the states of Wake, SWS and REM sleep. The mean and its corresponding 95 % confidence intervals are shown for each plot. **b** Power spectrum exponents calculated through ordinary least-squares fit on a log-log scale (OLS) or through the FOOOF parametrized spectra (FOOOF) [60] which only includes the aperiodic component.

**Table S3:**
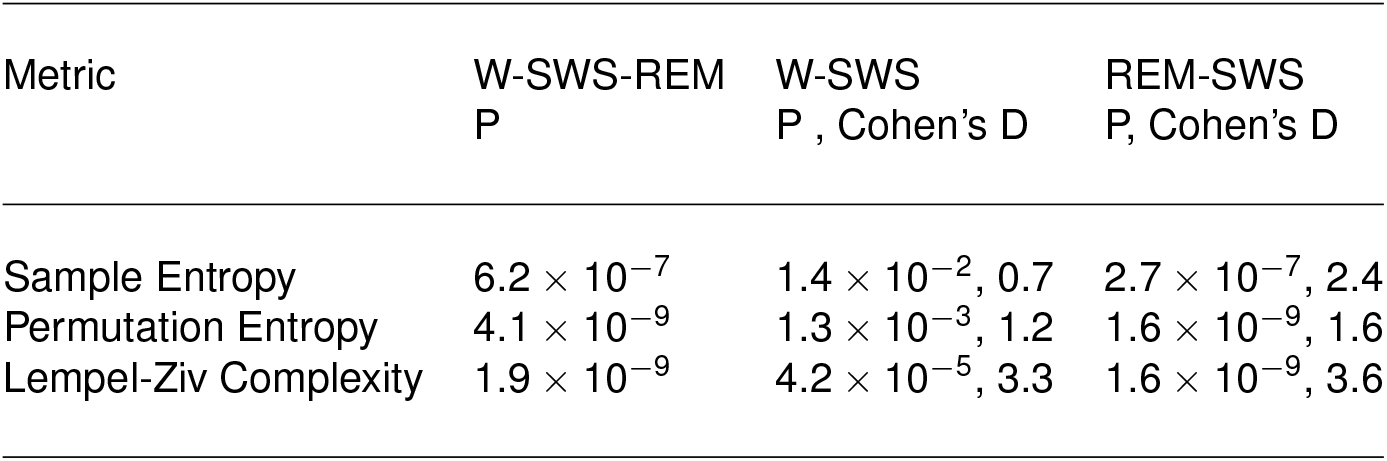
Statistical comparisons for the complexity metrics on real LFPs, shown in Fig 5d.

**Table S4:** Statistical comparisons for the complexity metrics on synthetic LFPs, shown in Fig 5d,e.

| Metric | W-SWS-REM<br>P | W-SWS<br>P , Cohen's D | REM-SWS<br>P, Cohen's D | SWS(All)-SWS(On)<br>P, Cohen's D |
| --- | --- | --- | --- | --- |
| Sample Entropy | $2.6 \times 10^{-7}$ | $3.0 \times 10^{-6}$, 1.35 | $3.0 \times 10^{-6}$, 1.5 | $1.6 \times 10^{-6}$, 1.6 |
| Permutation Entropy | $2.2 \times 10^{-7}$ | $1.0 \times 10^{-6}$, 0.4 | $1.1 \times 10^{-5}$, 0.8 | $1.1 \times 10^{-7}$, 1.2 |
| Lempel-Ziv Complexity | $2.5 \times 10^{-7}$ | $6.0 \times 10^{-6}$, 0.76 | $3.0 \times 10^{-6}$, 1.9 | $1.2 \times 10^{-7}$, 2.0 |

